# Learning High-Order Interactions for Polygenic Risk Prediction

**DOI:** 10.1101/2022.04.22.489134

**Authors:** Michela C. Massi, Nicola R. Franco, Andrea Manzoni, Anna Maria Paganoni, Hanla A. Park, Michael Hoffmeister, Hermann Brenner, Jenny Chang-Claude, Francesca Ieva, Paolo Zunino

## Abstract

Within the framework of precision medicine, the stratification of individual genetic susceptibility based on inherited DNA variation has paramount relevance. However, one of the most relevant pitfalls of traditional Polygenic Risk Scores (PRS) approaches is their inability to model complex high-order non-linear SNP-SNP interactions and their effect on the phenotype (e.g. epistasis). Indeed, they incur in a computational challenge as the number of possible interactions grows exponentially with the number of SNPs considered, affecting the statistical reliability of the model parameters as well. In this work, we address this issue by proposing a novel PRS approach, called High-order Interactions-aware Polygenic Risk Score (hiPRS), that incorporates high-order interactions in modeling polygenic risk. The latter combines an interaction search routine based on frequent itemsets mining and a novel interaction selection algorithm based on Mutual Information, to construct a simple and interpretable weighted model of user-specified dimensionality that can predict a given binary phenotype. Compared to traditional PRSs methods, hiPRS does not rely on GWAS summary statistics nor any external information. Moreover, hiPRS differs from Machine Learning-based approaches that can include complex interactions in that it provides a readable and interpretable model and it is able to control overfitting, even on small samples. In the present work we demonstrate through a comprehensive simulation study the superior performance of hiPRS w.r.t. state of the art methods, both in terms of scoring performance and interpretability of the resulting model. We also test hiPRS against small sample size, class imbalance and the presence of noise, showcasing its robustness to extreme experimental settings. Finally, we apply hiPRS to a case study on real data from DACHS cohort, defining an interaction-aware scoring model to predict mortality of stage II-III Colon-Rectal Cancer patients treated with oxaliplatin.

**Author summary:** In the precision medicine era, understanding how genetic variants affect the susceptibility to complex diseases is key, and great attention has been posed to Single Nucleotide Polymorphisms (SNPs) and their role in disease risk or clinical treatments outomes. Several approaches to quantify and model this impact have been proposed, called Polygenic Risk Scores (PRSs), but they traditionally do not account for possible interactions among SNPs. This is a significant drawback, as complex high-order SNP-SNP interactions can play an important role in determining the phenotype (a phenomenon called *epistasis*). Nevertheless, the number of possible combinations grows exponentially with the number of SNPs considered and including them in a predictive model becomes computationally challenging and affects the statistical reliability of the model. Some Machine Learning algorithms can answer this problem, but they are hardly interpretable. Here, we tackle these and other drawbacks of existing approaches proposing our novel PRS approach, *hi*PRS, that provides an interpretable weighted model with a user-defined number of predictive interactions. We designed it to handle typical real-life research scenarios, like small sample sizes and class imbalance, and we demonstrate here its superiority with respect to state-of-the-art methods.

## Introduction

Within the framework of precision medicine, it is becoming more and more important to stratify individual genetic susceptibility based on inherited DNA variation, an approach that progressed together with the latest advances in human genetics [1]. A substantial interest in leveraging the massive amount of genome-wide data now available has emerged, with the purpose of maximizing the predictive power of risk prediction models by incorporating the effect of Single Nucleotide Polymorphisms (SNPs) on the outcome [2]. Accurate genomic risk prediction has two great potential aims: to prospectively identify individuals at increased risk of disease, thus informing early interventions, and to aid diagnosis for diseases where current diagnostic approaches are imperfect [3].

One of the most traditional approaches to model genetic risk is the Polygenic Risk Score (PRS). PRS exploit a fixed model approach to sum the contribution of a set of risk alleles to a specific complex disease [4]. Calculating PRS is a common practice because of its simplicity, its computational efficiency, and the straightforward interpretability of the model itself. Indeed, while polygenic scores are used to predict phenotypes, there are other interests beyond forecasting. For instance, model interpretability is often key for research purposes such as the discovery or validation of SNPs’ role in disease risk. The schema in Fig.1 gives an overview on the strenghts and weaknesses of the most common PRS approaches in the literature.

**Fig 1.**
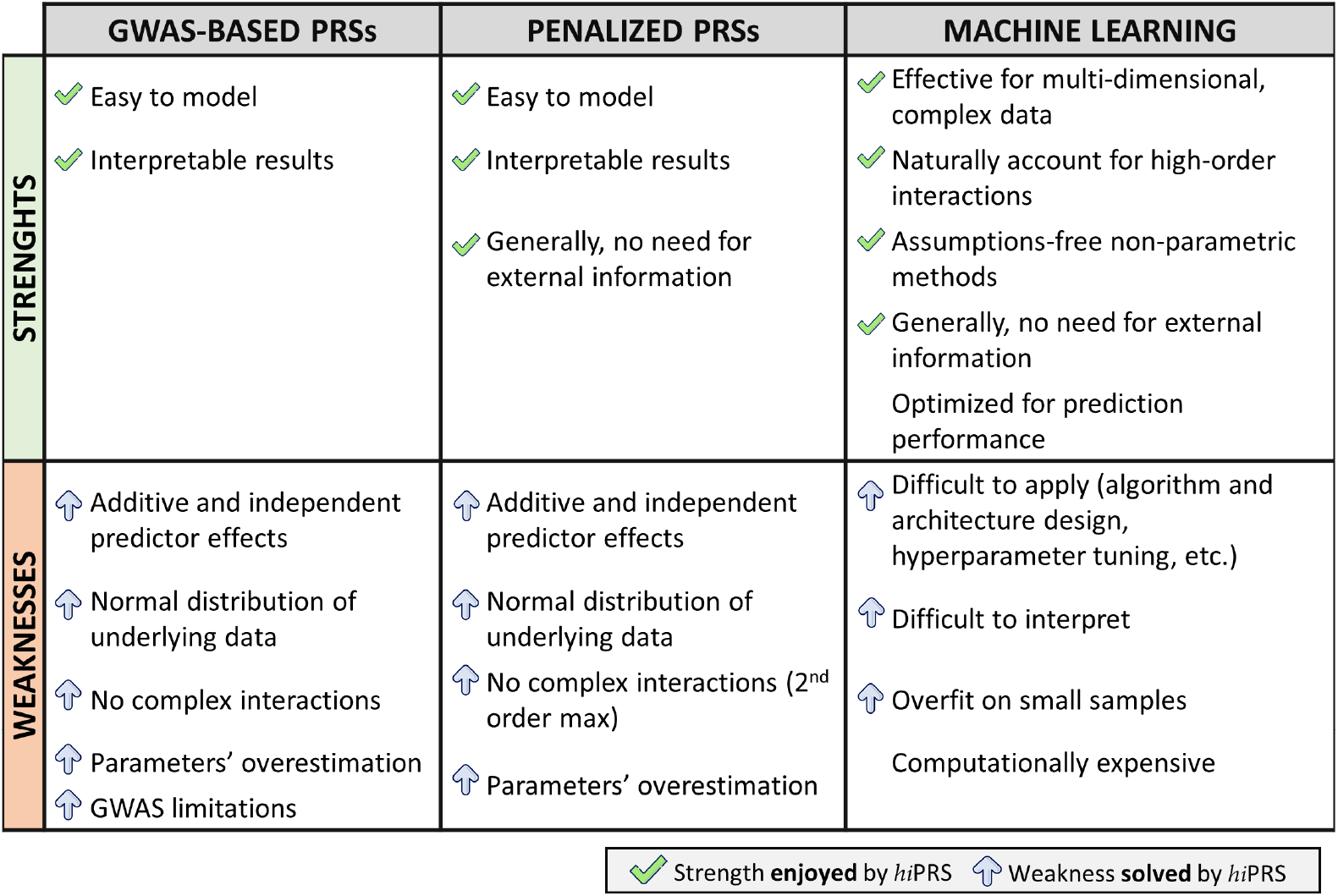
Alternative approaches to Polygenic Risk Scoring vs *hi*PRS. Strengths (first row) and Weaknesses (second row) of three main categories of PRS methods discussed in the Introduction. The green tick signals that the given point of strength applies to *hi*PRS as well. The blue arrow signals a point of weaknesses that *hi*PRS algorithm does not suffer, or some aspect that the algorithm was specifically designed to solve.

Standard weighted PRS estimation relies on Genome-Wide Association Study (GWAS) summary statistics obtained on one or more discovery cohorts modeling the independent effect of individual SNPs on the outcome. These PRSs (cf. Fig.1, first column, GWAS-based PRSs) exploit SNP-specific odds ratios or effect sizes to weight the contribution of the risk alleles on the disease risk or outcome [5, 6]. The set of SNPs to be included in this estimation may of course affect the score’s predictive power significantly. Some approaches include all SNPs, with the risk of incorporating useless or redundant information, while others retain a subset of SNPs based on predefined criteria (e.g., those passing an arbitrary p-value threshold in the GWAS results [5]).

Despite their wide adoption and appreciated simplicity and interpretability, the performance and reliability of traditional PRSs have been largely discussed and several methodological concerns have been raised (see, e.g., [7] and references therein) as the approach presents some evident limitations.

In particular, (i) the GWAS studies exploited for weights estimation are oftentimes underpowered by multiple independent testing and their effect sizes overestimated because of winners’ curse and biases [8–11]. Additionally, (ii) GWAS effect weights are traditionally computed considering the effect of single SNPs on the phenotype. By doing so, they do not account for the complex and nonlinear interactions between alleles in the genotype and their role in determining the phenotype, or the epistasis effect [4, 12–15]. Essentially, *epistasis* refers to departure from independence (or additivity, from a statistical point of view) of the effects of multiple loci in the way that they combine to cause disease [16]. In other words, interaction effects exist between loci, and their presence was reported to make major contributions to phenotypes [12, 13, 15, 17–20]. However, including SNP-SNP interactions in GWAS and risk scoring models is computationally challenging due to the high dimensions involved: in fact, the number of possible interactions grows exponentially with the number of SNPs considered. Additionally, this also increases dramatically the amount of independent tests to be carried out, thus strongly affecting their reliability. The authors in [21] attempted the inclusion of SNP-SNP interactions by filtering those that where relevant accordingly to some imposed biological criteria, but only managed to consider up to second-order interactions. Other methodological concerns of GWAS-based approaches are the fact that (iii) GWAS weights ignore the mediating role of clinical covariates when estimating SNPs effect on complex diseases [7], and that (iv) those models incorporate strict assumptions, e.g., they include additive and independent predictor effects, and assume that observations are uncorrelated [4, 14, 22]. These assumptions do not necessarily hold true when modeling complex polygenic diseases. For instance, Linkage Disequilibrium, i.e., the non-random association of alleles at two or more loci in a population [4], statistically translates into strong correlation between predictors. To account for LD some methods were developed to optimize SNPs *reweighting* (LD pruning and p-value thresholding, LDpred [23], and others).

Other well-recognized solutions to polygenic risk scoring discard GWAS weights and directly take genotype data as input, including various forms of penalization to restrict the pool of predictors (cf. Fig.1, Penalized PRSs). These include genomic BLUP [24], shrinkage methods (e.g. LASSO) or Generalized Linear Mixed Models (GLMMs). However, all these methods share with the most traditional PRSs the assumption of additive effect of single SNPs on the outcome, and may incur in the curse of dimensionality when trying to include all potential interaction terms, with the risk of overestimating effect sizes and obtaining unreliable models. This is particularly true if modeling high-order interactions. Nevertheless, these complex interactions were found useful in describing genotype-phenotype relationships for complex traits and common diseases both in humans and model organisms [25–29], highlighting the need for novel approaches able to account for high-order interactions.

In this respect, great attention has been recently devoted to polygenic risk prediction via Machine Learning (ML) algorithms (Fig.1, Machine Learning). These algorithms employ multivariate, non-parametric methods that robustly recognize patterns from non-normally distributed and strongly correlated data [15, 20, 22]. Moreover, these methods are naturally capable of modeling highly interactive complex data structures, making them powerful tools for complex disease prediction [15, 20, 22]. However, most ML algorithms, s.a. Random Forests (RFs), Support Vector Machines (SVMs) or Neural Networks (NNs), demonstrate great predictive power but lack in model interpretability, leaving researchers with no information on the role and structure of the complex interactions influencing the phenotype prediction. Moreover, these algorithms require a lot of training samples to avoid overfitting, but the available cohorts in real world research settings are oftentimes quite small.

In this work, we aim at addressing polygenic risk prediction proposing a novel PRS approach, called High-order Interactions-aware Polygenic Risk Score (*hi*PRS), whose most remarkable feature is the capability of robustly and reliably incorporating high-order interactions in modeling polygenic risk, while constructing a simple and interpretable model.

In brief (cf. Fig.2), *hi*PRS treats genotype-level data, and starts with a user pre-defined list of SNPs of interest. The algorithm exploits Frequent Itemset Mining (FIM) routines to build a list of candidate interactions of any desired order. This is achieved by scanning the observations in the positive class only, and by retaining those terms that have an empirical frequency above a given threshold *δ*. These candidates are subsequently ranked according to their relationship to the outcome in terms of Mutual Information (MI). From there, we select a restricted pool of *K* interactions to include in the final PRS, where *K* is user-specified. To this end, *hi*PRS incorporates a novel interaction selection algorithm inspired by the minimum Redundancy Maximum Relevance (mRMR) literature. More precisely, the algorithm selects *K* terms through the greedy optimization of the ratio between MI (*relevance*) and a suitable measure of similarity for interactions (*redundancy)*. This leads to a set of predictive, yet diverse, interactions that we use to define the score. In the end, the latter is built by weighting the contribution of each interaction term accordingly to the weights obtained when fitting a Logistic Regression (LR) model.

**Fig 2.**
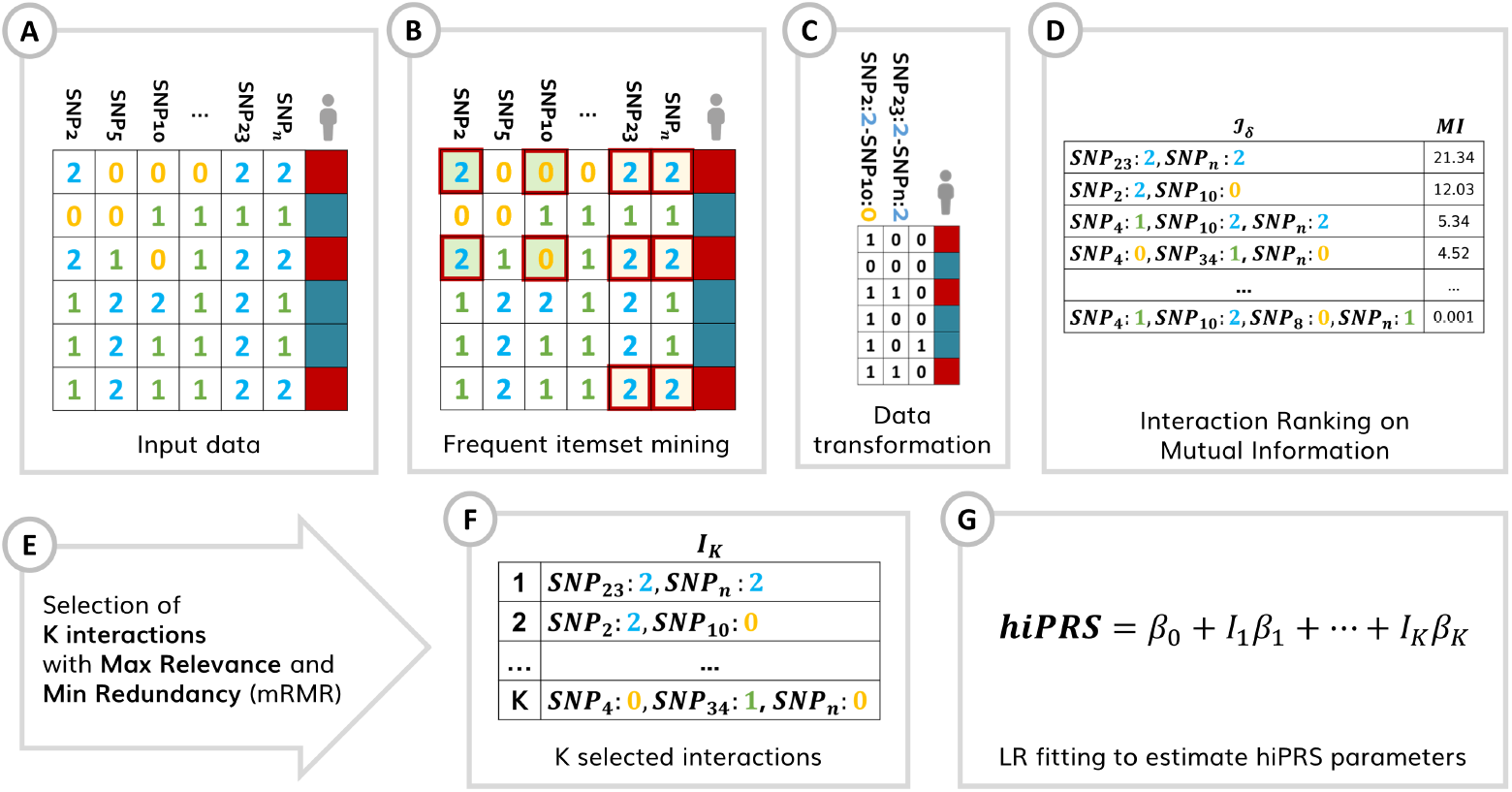
*hi*PRS Algorithm process flow. **(A)** Input data is a list of genotype-level SNPs. **(B)** Focusing on the positive class only, the algorithm exploits FIM (*apriori* algorithm) to build a list of candidate interactions of any desired order, retaining those that have an empirical frequency above a given threshold *δ*. This leads to a filtered set of terms in the form of sequences of pairs of SNP and associated categorical level (i.e., allele frequency in this example). The sequences can include from a single SNP-allele pair up to a maximum number of pairs defined by the user (*l*_max_). **(C)** The whole training data is then scanned, searching for these sequences and deriving a re-encoded dataset where interaction terms are binary features (i.e., 1 if sequence *i* is observed in *j*-th patient genotype, 0 otherwise). From this dataset we can compute the MI between each interaction and the outcome and **(D)** obtain a ranked list (*I*_*δ*_) based on this metric. Starting from the interaction at the top of *I*_*δ*_, *hi*PRS constructs *I*_*K*_, selecting *K* (where *K* is user-specified) terms through the greedy optimization of the ratio between MI (*relevance*) and a suitable measure of similarity for interactions (*redundancy)* (cf. Algorithm 1, Materials and Methods). This leads to a set of predictive, yet diverse, interactions that **(F)** we use to define the score weighting their contribution by fitting a LR model and retaining the corresponding *β* coefficients.

Similarly to traditional methods, *hi*PRS is based on a simple and interpretable weighted model. However, as highlighted in the summarizing scheme in Fig.1, our proposal overcomes the limitations of both traditional PRSs and modern Machine Learning approaches. In particular,

1. Compared to PRSs methods, *hi*PRS can include high-order interactions in the model without affecting weights estimation. Furthermore, it does not rely either on GWAS summary statistics nor on any other type of external information, and it does not suffer from the effect of predictors correlation.
2. *hi*PRS differs from most ML algorithms in that it provides an easily accessible, readable and interpretable model not meant for risk prediction only, but for discovery and validation of genetic susceptibility as well. Additionally, it is able to control overfitting, even on small samples.
3. On top of that, *hi*PRS is designed to be robust to class imbalance, which is oftentimes an issue when modeling rare complex diseases. Finally, we mention that, with very little effort, *hi*PRS can be generalized to include clinical covariates in the search for interactions, thus recovering their mediating role in determining SNPs effect on the outcome.

In the present work we demonstrate through a comprehensive simulation study the superior performance of *hi*PRS w.r.t. state of the art methods belonging either to Penalized PRSs or ML literature (cf. Fig.1), both in terms of scoring performance and interpretability of the resulting model. Moreover, we test *hi*PRS against complexities not seldom arising in real life research scenarios, such as small sample size, class imbalance and the presence of noise, showcasing its robustness to extreme experimental settings. Finally, we apply *hi*PRS to an interesting case study on real data from the DACHS cohort [30], defining an interaction-aware scoring model to predict mortality of stage II-III Colon-Rectal Cancer (CRC) patients treated with oxaliplatin.

## Results

### Simulation Study

This section describes the results of a large set of experiments run on simulated data whose generative mechanism was designed to present complex non-linear dependencies between a binary outcome *Y* and high-order SNP-SNP interactions. In particular, the positive class in the generated data, namely {*Y* = 1}, was defined in terms of the rules reported in Fig.3, which are ultimately high-order interactions. Additionally, each of the observed outcome values was mislabeled with probability *ε* (hereby called *random noise*), in an attempt to make the relationship between the outcome and the predictors less deterministic. More details on the generative algorithm can be found in Materials and Methods (cf. Simulated Data with Non-Linear Interaction Effect).

#### *hi*PRS outperforms benchmark algorithms in prediction performance

To judge the performance of *hi*PRS, we run a first experiment on simulated data and we compared the results with those of a comprehensive set of benchmark algorithms coming both from the traditional and more recent literature on polygenic risk prediction. Note that, to ensure a fair comparison, we only included methods that did not rely on any type of external information (e.g. summary statistics) besides individual-level genotype. More precisely, we picked methods from the following two classes: PRSs taking raw SNP values as input (Penalized PRSs) and algorithms from the ML literature. In the first group we included Lasso [31], Ridge [32], Elastic-Net [33] and glinternet [34]. The first three are penalized LR-based PRS methods with additive main effects and no interactions, while the last one is a penalized LR with group-Lasso regularization that includes second-order interactions. Conversely, as ML methods, we included two scoring algorithms accounting for high-order interactions: the approach proposed by Behravan et al. [35], that exploits XGboost for interaction selection and SVM for classification, and the one by Badre et al. [36], that relies on a Deep Neural Network (DNN) model. Specific technical details on each of these approaches, together with their implementation and chosen hyperparameters, can be found in the Materials and Methods section (cf. Benchmark Algorithms). All of this was considered within an experimental setting of mild complexity (i.e. reasonable sample size and low noise), consisting of 30 independent simulations.Indeed, for each of these we generated 1000 observations for training and 500 observations for testing. The mislabing *noise* probability was set to *ε* = 0.01.

The glinternet algorithm allows the user to define the number of interaction terms to include in the model, which we set to 3 and 8. Note that this method includes main effects of all considered interactions and estimates a parameter for each categorical level and their products. Therefore, 8 interaction terms actually correspond to more than 8 *×* 3^2^ = 72 regressors in the LR model (in particular, in our experiments we obtained 93 of them). In light of this, we chose 8 as maximum number of interactions, as that reflected the amount of generating rules in the simulated data (cf. Fig. 3), while the value of 3 is meant to test the performance of a simpler model. Differently from glinternet, in *hi*PRS the number of interaction terms *K* actually corresponds to the final dimension of the model. Here, we set *K* = 10, 40, resulting in two models of different complexity. In principle, both of them have enough degrees of freedom to discover the generative model, nevertheless, they both allow for handy readability and interpretability of the fitted model. We did not impose any limit on the order of the interactions, whereas we set the frequency threshold *δ* to 0.05.

**Fig 3.**
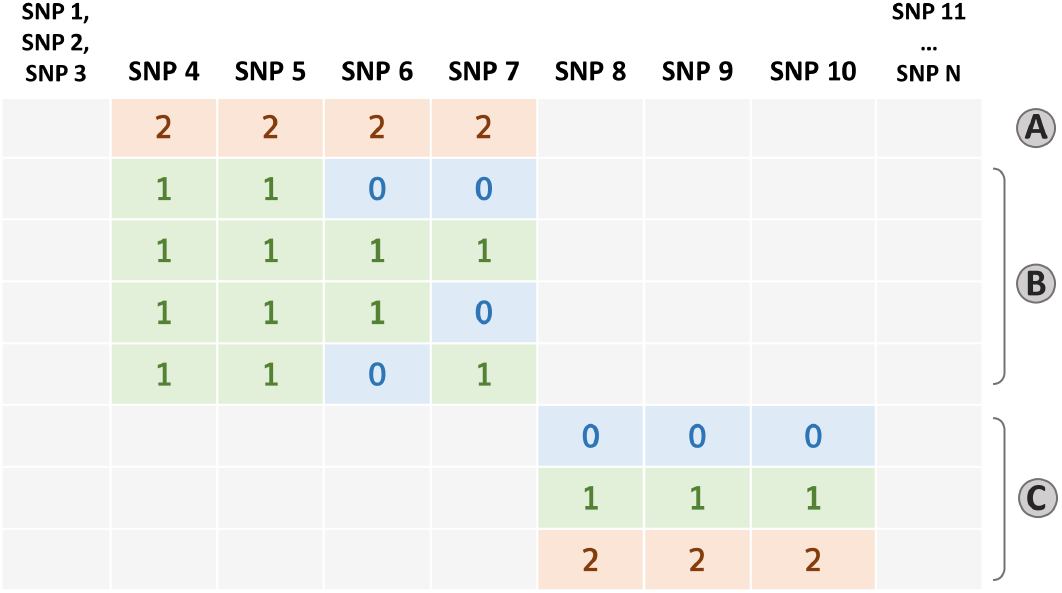
Simulation data generating rules. Graphical representation of the three rules that determine the positive class. Cases are obtained when **A B C**. More details on the generative model are provided in the Materials and Methods Section.

In Fig 4 we report the boxplots of the performance metrics in the 30 independent trials. The three Penalized PRSs behave similarly, with an average AUC around 0.6 and an Average Precision (AP) slightly above 0.3. Their performance reflects the maximum predictive power achievable by additive PRSs in the presence of epistatis. SVM-Behravan has the worst performance among the classifiers on both metrics, probably due to the fact that despite the SNP selection step accounts for interactions, it is followed by a linear SVM classifier that considers additive effects only. The DNN is unsurprisingly extremely performant, but its black box nature does not allow us to identify the role of the predictors in scoring observations. Moreover, whilst still high, it has a lower and much more variable AP performance. This might be due to the tendency to overfit the small sample of positive observations during training, possibly caused by the very large number of parameters typical of these models. Glinternet best AUC performance is achieved when including 8 interactions. However, the same model has a significant drop in AP, falling behind its simpler version with 36 terms: this goes to show that, despite the inclusion of more predictors, the identified interactions are not necessarily the most useful to identify the class of interest when the data is imbalanced. Moreover, the 93 parameters associated to the 8 interactions could easily overfit the underrepresented positive class. The best performer overall is *hi*PRS modeling 40 terms, showcasing the ability of our approach to identify predictive interactions that generalize well irrespectively of the under-representation of the positive class. The results obtained for *K* = 10 are also remarkable, especially if we consider that they correspond to the model having the least number of parameters.

**Fig 4.**
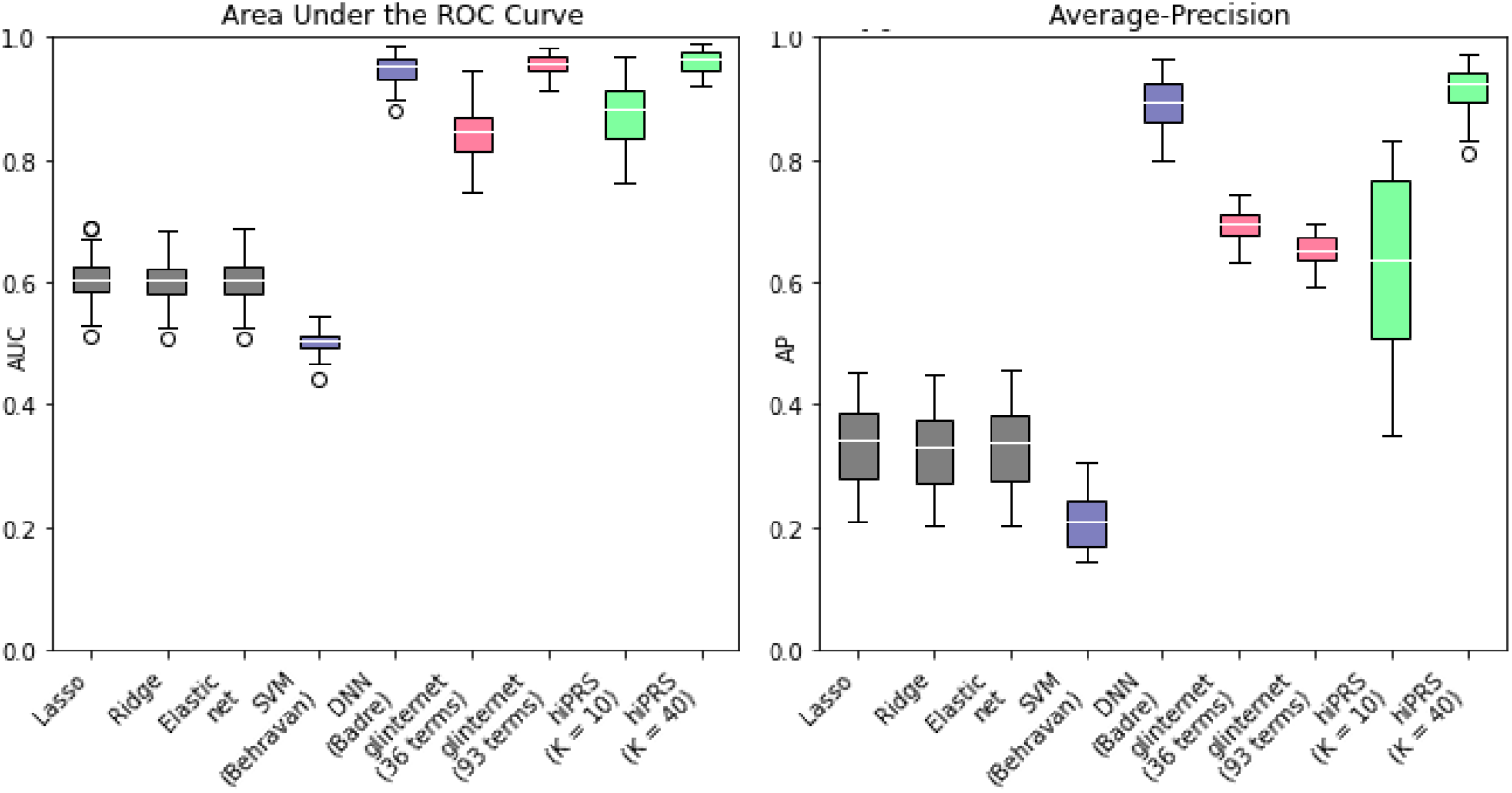
*hi*PRS Results on risk prediction against benchmark PRSs and ML approaches. AUC (left) and AP (right) performance distributions of 30 independent trials. In grey the three traditional penalized PRSs approaches with additive effects only; in violet the two ML algorithms (SVM-Behravan and DNN-Badre); in pink glinternet algorithm for two model dimensions (3 interactions, i.e. 36 terms, and 8 interactions, i.e. 93 terms); in green *hi*PRS for *K* = 10 and *K* = 40.

#### *hi*PRS captures and explains interaction-based generative mechanisms better than benchmarks

One of the added values of *hi*PRS is the capability of capturing dependencies among predictors, by selecting the most relevant to determine the phenotype and presenting them within a simple and interpretable model. In many research settings, a slight loss in prediction quality may be acceptable if it leads to a more meaningful interpretation of the predictors [37, 38]. To test the ability of *hi*PRS in capturing them, we focused on one of the previously mentioned datasets, and checked the 10 interactions selected by the simplest model, *K* = 10. We report the selected interactions in Fig.5. We mention that, to ensure a fair comparison, we picked the dataset where the worst performer, SVM-Behravan, achieved one of its highest AUC. This dataset contained 182 cases in the training set, i.e. 18.2% of the total. The bar chart in Fig.5 reports the absolute frequency of the generating rules available in the positive class.

*hi*PRS captures most of rule C, assigning very high positive betas to the first, second and fourth interaction. Combined together, the fourth and fifth selected interactions partially recover rule A, while the third one almost fully captures rule B.3. Note that, among positive samples in the training data, B.3 is the most frequent version of rule B (see Fig. 5, left panel). It is also interesting to note that *hi*PRS finds two protective terms, namely *{*SNP_4_ = 0} and *{*SNP_5_ = 0}, that both nullify rules A and B.

**Fig 5.**
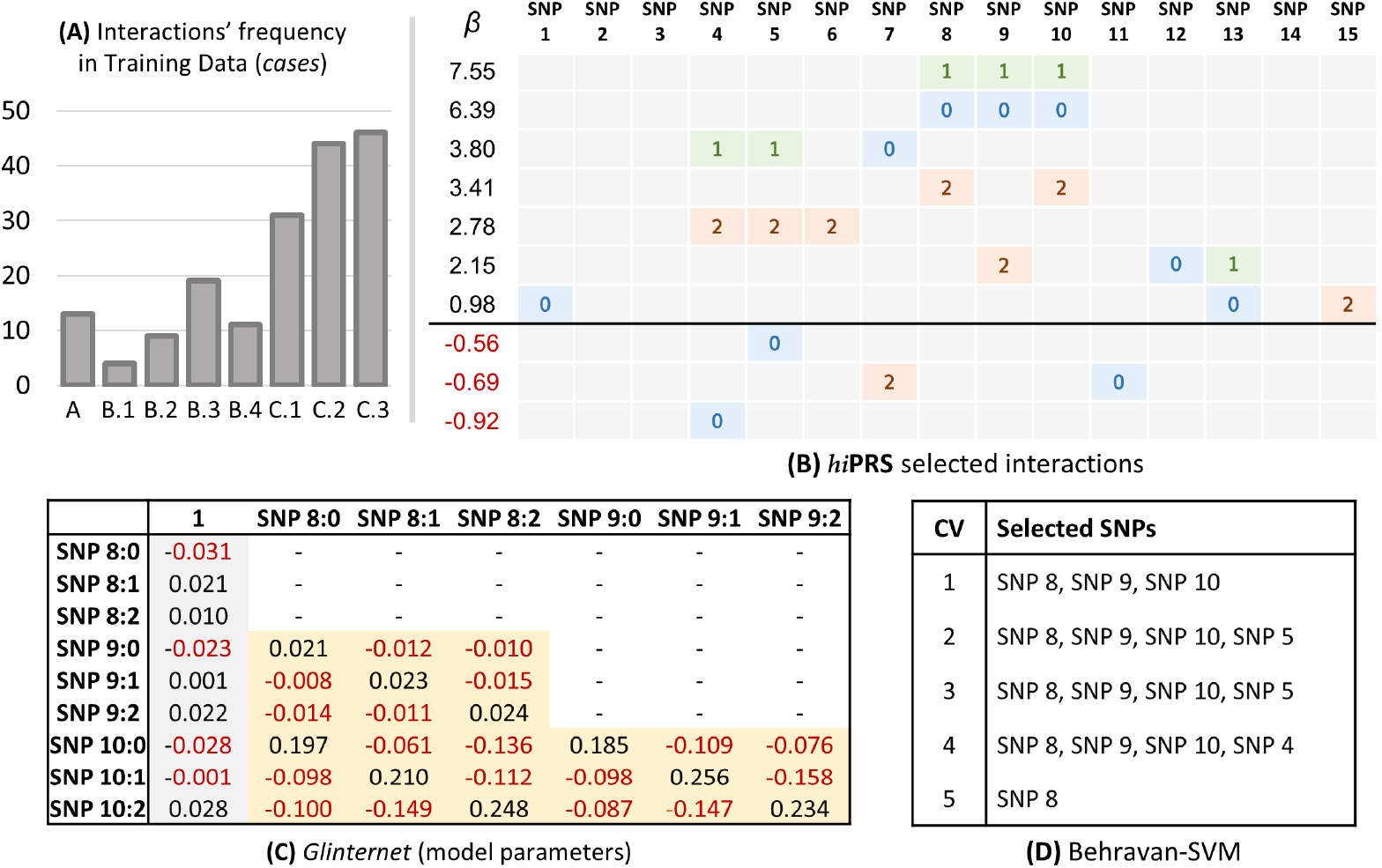
Interpretability Analysis. (A) Absolute frequency of the generative rules in the training data, limited to the positive class. (B) Interactions selected by *hi*PRS with *K* = 10 and corresponding *β* coefficients. (C) Coefficients of the glinternet model with 3 interaction terms: main effects are in gray, interactions in yellow. (D) Lists of SNPs selected by SVM-Behravan during its five internal cross validations, cf. Benchmark Algorithms in the Materials and Methods Section. Note: reported results are limited to one simulation among the 30 randomly generated datasets.

Let us now briefly discuss the results obtained by the competitors. In panel (c) of Fig. 5 we report the model parameters for glinternet when modeling 3 interactions. Here, we see that the good performance of glinternet is granted by the inclusion of three interaction terms in the model, which, despite being of second order only, are subsets of the true generating rule C. Moreover, the estimated effect sizes are correctly positive for couples of identical allele frequencies only (i.e., the diagonal of the matrices of coefficients associated to each interaction). However, due to extremely large number of parameters, it is very hard to inspect the model and drive suitable conclusions.

Furthermore, main effect sizes are not truly relevant in determining the phenotype and the generative mechanism is only captured partially, as rule C is the only one that is actually identified. Within the same Figure, but in panel (d), we list the SNPs selected by SVM-Behravan during its 5-fold cross-validation. Notably, SVM-Behravan is able to recover some of the SNPs associated with the generating rules, however, we are left with no information on the structure of the interactions.

#### *hi*PRS can deal with extremely small sample size

We tested *hi*PRS performance for small sample sizes. We believe this to be a relevant setting, as in most real research scenarios clinicians have to deal with individual-level data of significantly small cohorts. To this end, starting from the sample size that we considered in our previous experiments, we repeated our analysis on a sequence of four decreasing sample sizes, namely *n* = 1000, 750, 500, 250. That is, for each *n*, we run 30 independent simulations, and we registered *hi*PRS performance in terms of AUC and AP. To ensure comparable results, the noise level was fixed to *ε* = 0.01 for all experiments, and the model was always evaluated on a test set of 500 instances. Once again, we tested *hi*PRS with *K* = *{*10, 40}, while *δ* = 0.05 as before. Results are reported in Fig 6, panel A. Despite the unavoidable decrease in performance, we note that even for very small samples, 250 observations, *hi*PRS is able to provide insightful results, with AUC levels of 0.8 and *∼* 0.7 respectively. AP is lower, *∼* 0.5 for *K* = 40, but still significantly better than traditional penalized PRSs when trained on 1000 observations. Nonetheless, note that a training sample of 250 observations in total corresponds to less then 75 cases to learn from (cf. Simulated Data in Materials and Methods), which is an extremely challenging setting. For 500 training samples (i.e. *∼* 150 cases at most) *hi*PRS with *K* = 40 reaches almost 0.9 AUC and an AP of *∼* 0.7. 260 These results testify in favour of *hi*PRS generalization potential, which is likely induced by the mRMR-based interaction selection algorithm. Indeed, the latter is optimized to select the most predictive features, while favouring the introduction of *diverse* information in the model. Minimizing redundancy in the selection makes *hi*PRS less prone to overfitting, irrespectively of the sample size or the number of cases.

**Fig 6.**
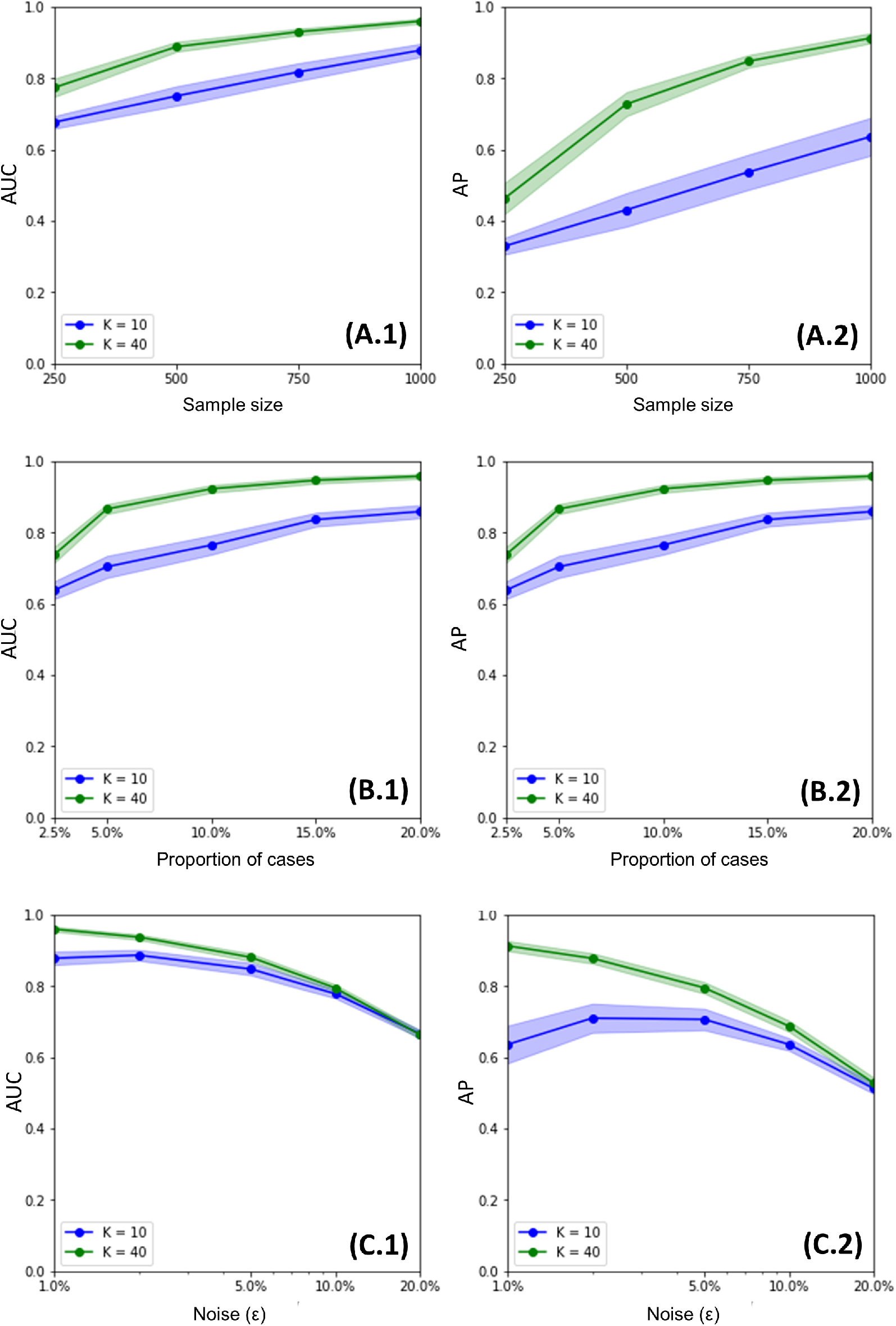
Sensitivity Analysis results. Average performance of *hi*PRS in terms of AUC and AP for variable sample size (A.1 and A.2), class imbalance (B.1 and B.2) and missing heritability, i.e. noise (C.1 and C.2). Confidence bands are at the 95% level. The *x*-axis is in logarithmic scale for panels C.1 and C.2.

#### *hi*PRS is robust to class imbalance

To test *hi*PRS against extreme class imbalance, we had to modify slightly our procedure in the generation of the simulated data. This is because although our generative mechanism is based upon random generation of allele categories and random noise (*ε*), it also relies on a deterministic rule-based definition of the positive class. Therefore, the positive class will always appear in the data with an approximately fixed frequency depending on *ε* (cf. Materials and Methods). We overcame this drawback via undersampling. More precisely, let 0 *≤ q ≤* 1 be any wished proportion. To obtain a training sample of 1000 instances and 1000*q* cases, we generate a larger dataset where observations are added iteratively until there are at least 1000*q* cases and 1000(1 *− q*) controls, then we discard all exceeding observations. We adopted this procedure for a varying proportion of cases, namely *q* = 2.5%, 5%, 10%, 15% and 20%. For each of those values, we generated 30 independent training sets, and measured *hi*PRS performance over as many independent test sets of 500 instances. We mention that, while we used subsampling to generate the training data, we stuck to our usual approach for the test data. Indeed, the complexity lies in training models on imbalanced data; furthermore, the distribution of classes in the test set has no impact on the metrics that we use for evaluation, i.e. AUC and AP.

In Fig. 6, panel B, we report the results for this simulation study. Notably, even for the smallest proportion of observations in the positive class (2.5%), *hi*PRS succeeds in learning a sufficiently generalizable set of interactions, with both average metrics above 0.6 when *K* = 10, and above 0.75 when *K* = 40. Moreover, the larger model approaches 0.9 on both AUC and AP for a proportion of cases of 5%, meaning only 50 observations to learn from.

#### *hi*PRS is robust to variability (missing heritability)

Missing heritability is the proportion of variance in the phenotype that cannot be explained by genotype information [39]. This variability can be induced by several factors [39]: one is the need to account for SNP-SNP interactions when modeling phenotypic traits [40], but it can be nonetheless induced by factors that cannot be modeled with the data at hand. In our simulation setting, we reproduce the effect of missing heritability via the *noise* parameter, as the two can be easily linked together. The explicit way by which the missing heritability depends on *ε* is detailed in Materials and Methods Section, equation (2). To test *hi*PRS robustness to variability in the training data, we generate 30 training and test set, respectively with 1000 and 500 observations, for each of the following noise levels, *ε* = 0.01, 0.02, 0.05, 0.1 and 0.2. Results are in Fig 6, panel C. Note that, up to a noise level of 10% (i.e. *∼* 100 mislabeled observations in the training set, and *∼* 50 in the test set), both models maintain an average AUC near 0.8. This is considerably interesting if we consider that a noise of *ε* = 0.1 corresponds to a missing heritability of 51%, i.e. a setting where the SNPs themselves can only explain at most 49% of the target variability.

### Case Study on Real Genotype Data

Oxaliplatin-based chemotherapy is a standard treatment to treat colorectal cancer patients, including stage III and selected stage II patients. However, response rates to oxaliplatin vary substantially between patients and some patients suffer from serious side effects caused by the treatment. Therefore, additional consideration of genetic markers to identify patients who are more likely to benefit from oxaliplatin would be highly desirable [41]. Here, we exploited *hi*PRS to build a polygenic risk score that could predict the short-term survival of oxaliplatin treated patients. The analysis included 349 stage II-III patients who received oxaliplatin from an ongoing population-based case-control study (DACHS, colorectal cancer: chances for prevention through screening). We refer to the Material and Methods Section for a detailed description of the data. We restricted our attention to those patients that either survived (0) or died within three years (1), and we constructed the PRS on the base of nine recently validated SNPs.

To account for the small sample size, the analysis was carried out via cross-validation on 4 folds. This means that the whole procedure required to build the *hi*PRS was repeated on each fold separately, with a common choice of the hyperparameters *δ* and *K*. The values of the two were optimized via grid-search and resulted in *δ* = 15.5% and *K* = 5. To highlight the improved performance derived from the inclusion of high-order interactions in the model, a Penalized PRS with Lasso regularization (i.e. modeling additively SNP’s main effects only) was also implemented.

Results are shown in Fig.7. The boxplots report the AUC values obtained by the two PRSs across the four folds of the cross-validation procedure. Despite the small number of training data, it appears that *hi*PRS is able to find interactions that generalize well and that are far more predictive with respect to single SNPs. Indeed, *hi*PRS reports an average AUC of 0.72 and uses as little as 6 terms, counting the intercept. In contrast, the Penalized PRS introduces 9 *×* 3 + 1 = 28 parameters but only manages to obtain an average AUC of 0.57. This result supports the idea that SNPs interactions may play a role in quantifying the response to oxaliplatin treatments.

**Fig 7.**
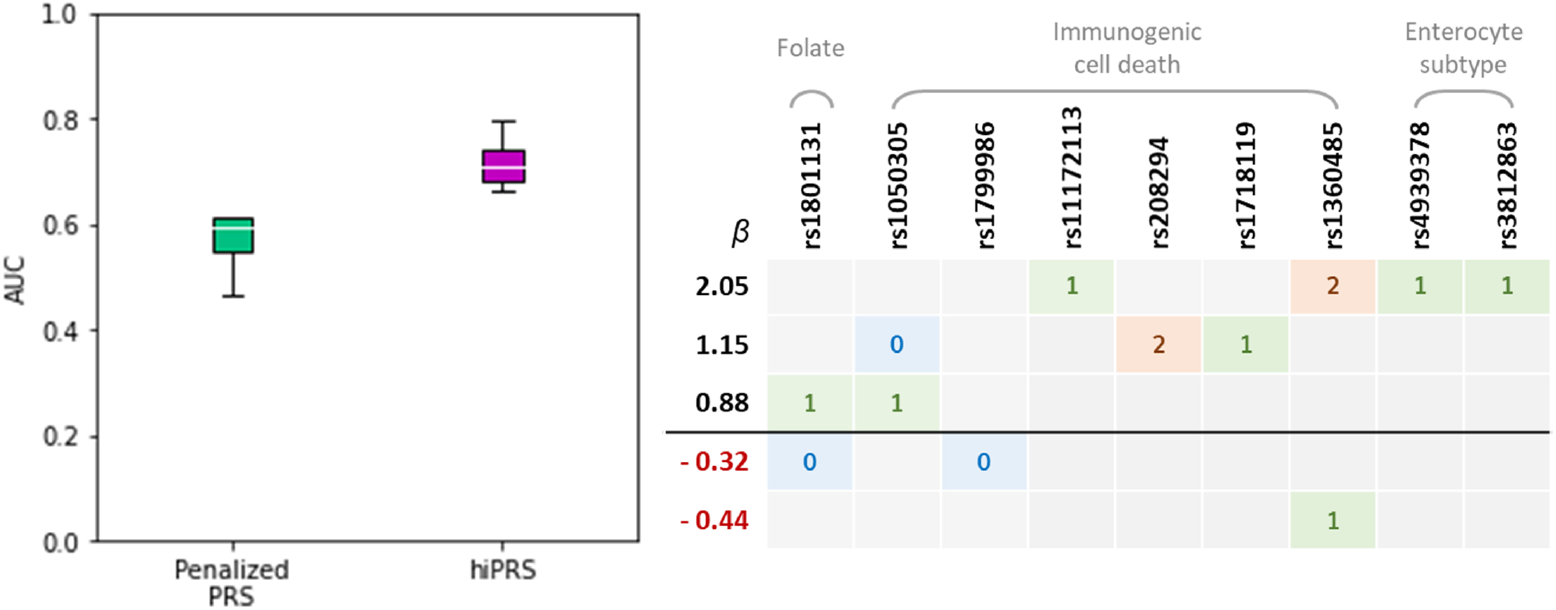
DACHS case study results. Left panel: AUCs obtained by *hi*PRS and a benchmark model during cross-validation (four folds). Average AUCs are 0.72 and 0.57 respectively for *hi*PRS and the benchmark model. Right panel: interactions selected by *hi*PRS, and corresponding effect-sizes, when fitting the model on the whole dataset. In grey are reported the pathways each SNP belongs to (cf. Materials and Methods).

At this regard, it is of interest to know which interactions were selected by the *hi*PRS routine. We have reported in Fig.7, right panel, the five interactions identified when fitting *hi*PRS on the whole dataset. Of note, we mention that one of the longest interactions, namely

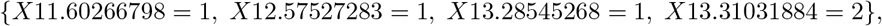

was also the most robust. Indeed, that was the only interaction to be always selected during the cross-validation routine.

## Discussion

In this work we presented *hi*PRS, a novel approach to tackle polygenic risk scoring that captures and models the effect on the phenotype of single SNPs and SNP-SNP interactions of potentially very high order. The algorithm takes individual-level genotype data as an input, overcoming potential biases of GWAS information and can work on any predefined set of SNPs to provide an easily interpretable tool not only for accurate prediction, but also to perform solid inference and SNP-SNP interaction discovery or validation. *hi*PRS allows the user to define the size of the model and the maximum order of the interactions to search for, which allows for reliable parameter estimations, especially for small samples, and convenient inspection by domain practitioners.

We have tested *hi*PRS against similar benchmark methods that rely on individual-level data, demonstrating its superior performance with respect to traditional PRSs and more complex ML-based methods. Indeed, *hi*PRS performs significantly better than any model accounting for additive effects only (i.e. Lasso, Ridge, Elastic-Net and SVM-Behravan) when the true generative mechanism incorporates SNP-SNP interactions, i.e., in the presence of *epistatic* effects. Moreover, its results are on pair with state-of-the-art methods accounting for interactions, such as glinternet, a more advanced penalized PRSs, and DNN-Badre, a nonlinear model based on artificial neural networks. Nevertheless, with respect to glinternet, *hi*PRS allows to include interactions of higher order with a much smaller set of parameters to estimate. ML-based approaches like DNN-Badre can instead model interactions of extremely high order and perform very well in patients’ scoring, as shown in our benchmark experiment, but tracing back the effect of all predictors or reconstructing the fundamental generating interactions from those complex black-box models is almost impossible. Even by applying to these models the wide variety of Explanation algorithms available, such as SHAP or LIME, one may at most quantify the role of each covariate alone, without however being able to retrieve any information on their interaction patterns.

There are some relevant facts about *hi*PRS that are worth mentioning. One lies in the fact that the algorithm explicitly defines and selects interaction terms as sequences of feature-level pairs. This aspect allows to gather fundamental insights on the true generating mechanism of the phenotype, especially in the presence of epistatic effects. For instance, there may be situations in which the same SNPs can have either a risk or a protective effect on the phenotype depending on the alleles’ configuration, as in the simulated example for which we reported *hi*PRS selected interactions (cf. SNP_4_ and SNP_5_ in Fig.5). In that case, only *hi*PRS consistently assigned a proper effect to each level of the two features, by estimating a coefficient for each interaction that included one of the predictive SNP-allele pairs. While a method like glinternet might capture the effect of specific genotypes, this comes at the expense of estimating a *β* parameter for each of the possible SNP-allele combinations. Conversely, *hi*PRS can directly select the predictive patterns only, and estimate their unique effect on the disease.

Besides all the relevant aspects mentioned above, in the present work we wished to demonstrate *hi*PRS’ capability to tackle the complexities of real research scenarios. To this end, we validated our method through an extensive set of simulations, showcasing its ability to deal with noise, strong class imbalance and even extremely small sample sizes without overfitting on the training data. This result was made possible thanks to the combination of the frequency threshold *δ*, which reduces the number of candidate interactions to those appearing in a sufficiently large portion of the positive class, and the mRMR-based interaction selection algorithm. Indeed, minimizing redundancy (while maximizing relevance) forces *hi*PRS to select the most diverse predictive interactions, which allows the model to generalize and capture the broader picture of the generative mechanism. This aspect was confirmed inspecting the 10 interactions selected in Fig. 5. While all competitors picked the SNPs defining the most frequent rules in training data only (i.e. rule C in that example), *hi*PRS managed to include instances of other less frequent generative rules, and assign these terms a large positive coefficient.

Besides testing *hi*PRS on simulated settings, we applied the algorithm to a Case Study on real genotype data. Results show a significant improvement in performance w.r.t. a traditional Penalized PRS. Notably, the two terms in the resulting model (cf. Fig.7, right panel) with the highest positive effect on the phenotype are third- and second-order interactions, which would require an extremely large number of predictors for their effect to be captured by traditional PRSs. Conversely, *hi*PRS estimates their effect together with other three terms only, allowing for an accessible inspection of the lightweight resulting model. The results presented in this case study were only validated internally via cross-validation, as further investigations were not feasible due to the lack of external data. Nevertheless, the experiment was meant to showcase the potential of *hi*PRS and its applicability on real data. Since the obtained results are promising, we plan to deepen the clinical interpretation and to validate the discoveries in future works, as soon as a comparable external dataset is available.

At the moment, one major limitation of *hi*PRS lies in the scalability of the algorithm, especially when the number of SNPs grows dramatically. In fact, the search of high-order interactions is intrinsically expensive, and the computational cost of the mRMR selection routine grows quadratically with the number of candidate interactions. For this reason, our algorithm is better suited for those contexts where the number of SNPs is limited, as in clinical studies that start from literature-validated SNPs. There, the computational cost can be almost negligible: for instance, fitting *hi*PRS always took less than 5 seconds in our experiments.

Nevertheless, the computational cost of *hi*PRS can be reduced in several ways, e.g. by increasing the frequency threshold *δ* or decreasing the maximum interaction length *l*_max_. Furthermore, *hi*PRS can easily accommodate modern approaches to Frequent Itemset Mining, such as GPU-accelerated versions of *Apriori* [42] or single-scan algorithms [43, 44], to boost the preliminary search of interactions.

## Conclusions

In conclusion, in this work we presented *hi*PRS and demonstrated how this simple yet effective novel PRS approach can represent a valuable tool for the analysis of genotype data in a precision medicine framework. Indeed, it can effectively be exploited in real life experimental settings with potentially very small sample sizes to model either common or rare phenotypes with the inclusion of high-order SNP-SNP interactions effects. We largely discussed potentials and limitations of our proposed approach, but another remarkable aspect to note, is that the method can flexibly accomodate any kind of categorical data, even though *hi*PRS was presented here to process SNPs data with allele frequency categories. For instance, it might be effortlessly applied to data describing genetic variants or epigenetic mutations encoded as binary variables, single and multi-level categorical clinical information, or their combination. This makes the *hi*PRS approach particularly interesting to tackle multi-omic studies, or to model the recognized interactions between genotype and environmental factors, or to investigate the mediating role of clinical conditions in determining the genotype effect on complex traits, widening the scope of applicability and relevance of our proposal.

## Materials and Methods

### Simulated Data with Non-Linear Interaction Effect

The synthetic data was built as follows. We considered a pool of *p* = 15 independent SNPs, *S*_1_,…,*S*_*p*_, and a binary outcome variable *Y*. In order to impose a nonlinear dependence between the two, we considered a model of the following form

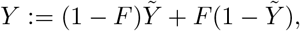

where 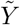 is univocally determined by the fifteen SNPs via the relation below,

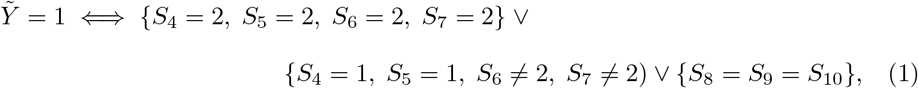

whereas *F ∼ B*(*ε*) is a binary random variable, independent on the SNPs, that we use to model either noise in the data or unexplained variability. Note in fact that if *F* = 0, then 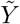 and *Y* coincide, otherwise the labels are flipped. It is worth noting the following facts.

1. Let 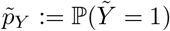 and *pY* := ℙ(*Y* = 1). It is easy to see that

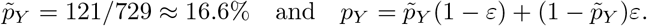

In particular, for *ε ≤* 0.2 we have *p*_*Y*_ *≤* 30%, resulting in a class imbalance where cases are the minority.
2. We can explicitely quantify the amount of variability in *Y* that cannot be explained by the SNPs only. Indeed,

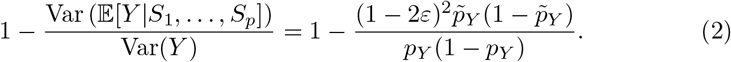

We refer to the above as *missing heritability*. In our final experiment, we use (2) to provide better insights on the *hi*PRS performance for different values of *ε*.

### DACHS Data

For our case study we considered genotype-level data and clinical information coming from an ongoing population-based case-control study (DACHS, colorectal cancer: chances for prevention through screening). Details of the study have been previously described by Brenner et al. in [30, 45]. The dataset originally included patients recruited between 2003 and 2015. Genotype and complete follow-up data (for either 3, 5, or 10 years) were available for a total of 3689 histologically confirmed cases diagnosed between 2003 and 2014.

We excluded patients who have metastatic disease, who had not received adjuvant chemotherapy, received neoadjuvant chemotherapy, had an unknown start date of chemotherapy, or died within 30 days of the start of chemotherapy. Of patients treated with first-line adjuvant chemotherapy, we included stage II-III patients who received at least four cycles of oxaliplatin-based treatments. Finally, in order to study the short-term survival, we excluded all patients that died after 3 years. This resulted in a subsample of 349 patients. The main characteristics of the study population are shown in Table 1. Details about genotyping and imputation for the DACHS population have been described in detail somewhere else, see [41]. Among the available SNPs, we focused on the nine reported in Table 2. These are of particular interest as they were only recently validated as associated with the efficacy of oxaliplatin-based treatment in CRC patients [41].

**Table 1.** Patient characteristics of the subsample analysed in the DACHS case study.

|  | Number (%) |
| --- | --- |
| Age (years $\leq 50$) | 52 (14.9%) |
| Sex (Female) | 130 (37.2%) |
| Stage (Stage II) | 28 (8.0%) |
| Grade (grade 3-4) | 109 (31.9%) |
| CRC site (Rectum) | 87 (24.9%) |
| Death | 50 (14.3%) |

**Table 2.**
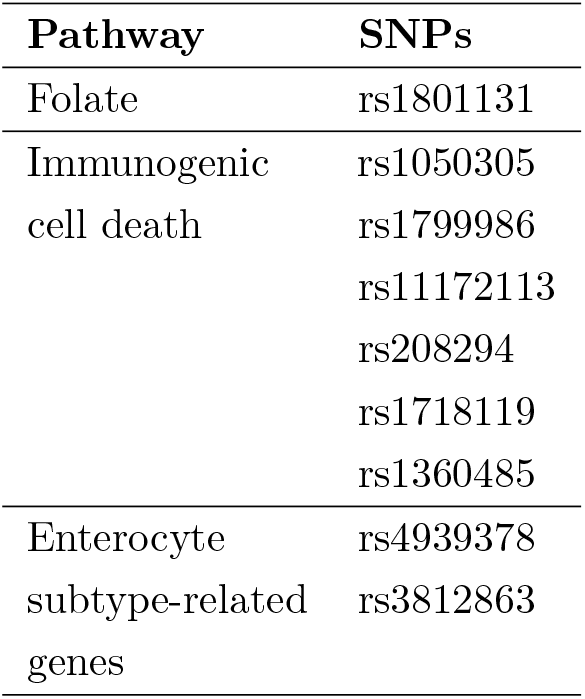
SNPs considered in the DACHS case study.

### Ethics Statement

The DACHS study was approved by the ethics committees of the Medical Faculty of the University of Heidelberg and the State Medical Boards of Baden-Wuerttemberg and Rhineland-Palatinate.

## Proposed methodology

### Preliminaries and Problem Statement

In order to describe our approach, we first introduce some notation. Let *S*_1_, …, *S*_*p*_ be *p* SNPs and let *Y* be a random variable denoting a binary outcome. We consider each SNP to be a categorical variable taking values in {0, 1, 2}, where labels read as follows. We denote major allele homozygosis by 0, heterozygosis by 1, and minor allele homozygosis by 2. To model SNP-alleles interactions, we use products of indicator functions. For instance, *I* := **1**_*{*0}_(*S*_1_)**1**_*{*1}_(*S*_2_) encodes the interaction between major allele homozigosis in *S*_1_ and heterozygosis in *S*_2_. We may then define the collection of all SNP-allele interactions as

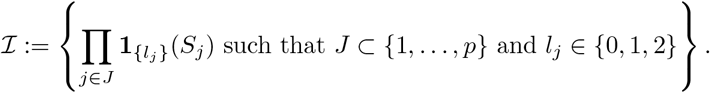

We note that, with little abuse of notation, the set *I* also contains the dummy-variables associated to the SNP-alleles, as those are obtained when *J* is a singleton.

We are given a dataset 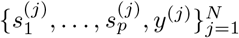 consisting of *N* i.i.d. realizations of the SNPs and the outcome *Y*. Starting from these, we aim to construct a PRS of the form

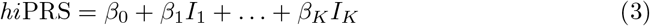

Where 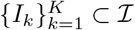 To this end, we propose a novel scoring method, *hi*PRS, where *K* is user-specified and a data-driven algorithm returns the list of interactions with the corresponding weights. We detail the whole idea in the following section.

### *hi*PRS Algorithm

For any *I* ∈ ℐ, let 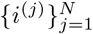 be the corresponding observations in the dataset. We define the collection of all cases and its complement as

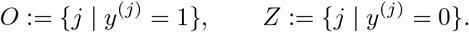

We make the assumption that |*O*| ≪ |*Z*|, which is the typical scenario of a rare outcome. We refer to *O* and *Z* respectively as the minority and majority class. As a first step, we scan the data relative to the minority class, and we search for those interactions 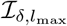 that appear with an empirical frequency above a given threshold *δ >* 0, and have length at most *l*_max_. More precisely, we define

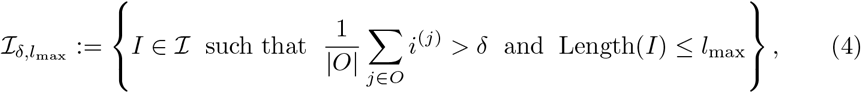

where the Length(*I*) is the number of SNPs involved in the definition of *I*. In principle, computing 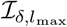 can be very demanding since | *ℐ*| = 3^*p*^. However, this drawback is mitigated by two key factors. First of all, we note that each interaction uniquely corresponds to a *pattern of alleles*. For instance, let

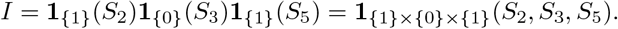

Then *i*^(*j*)^ = 1 if and only if the pattern *{S*_2_ = 1, *S*_3_ = 0, *S*_5_ = 0} is observed in the *j*th patient. This duality between interactions and patterns allows us to reframe (4) in the context of frequent itemsets mining, where we can relay on a moltitude of algorithms such as *Apriori* and *FP-Growth*. Additionally, the computational cost is alleviated by the fact that we limit the search of candidates to the minority class *O*.

The next step is to extract a suitable sublist 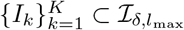 to be used in (3). While this problem can be framed in the context of feature selection, finding an optimal solution can be very hard due to the large number of candidates. To overcome this drawback, we introduce a filtering technique based on the so-called Minimum Redundancy – Maximum Relevance approaches, mRMR for short. The idea goes as follows. First, we introduce a relevance measure based on the (empirical) mutual information, that is

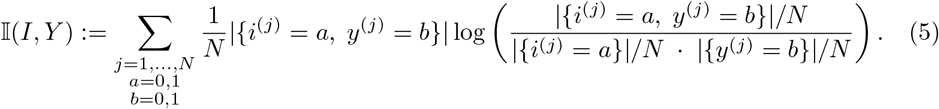

By definition, 𝕀 (*I, Y*) ≥ 0 and larger values are obtained when *I* and *Y* are informative with respect to each other. Parallel to this, we introduce a redudancy measure 𝕊 : ℐ × ℐ → ℕ that quantifies the similarity between two given interactions. More precisely, we define 𝕊 (*I, T*) to be the number of common alleles within the two patterns. We then construct the set 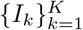 along the lines of mRMR methods, that is through the greedy optimization of the ratio between relevance and redundancy. We report a detailed pseudocode in Algorithm 1.

#### Algorithm 1

Extraction of the hiPRS allele-patterns, starting from a list of candidates and raw data.

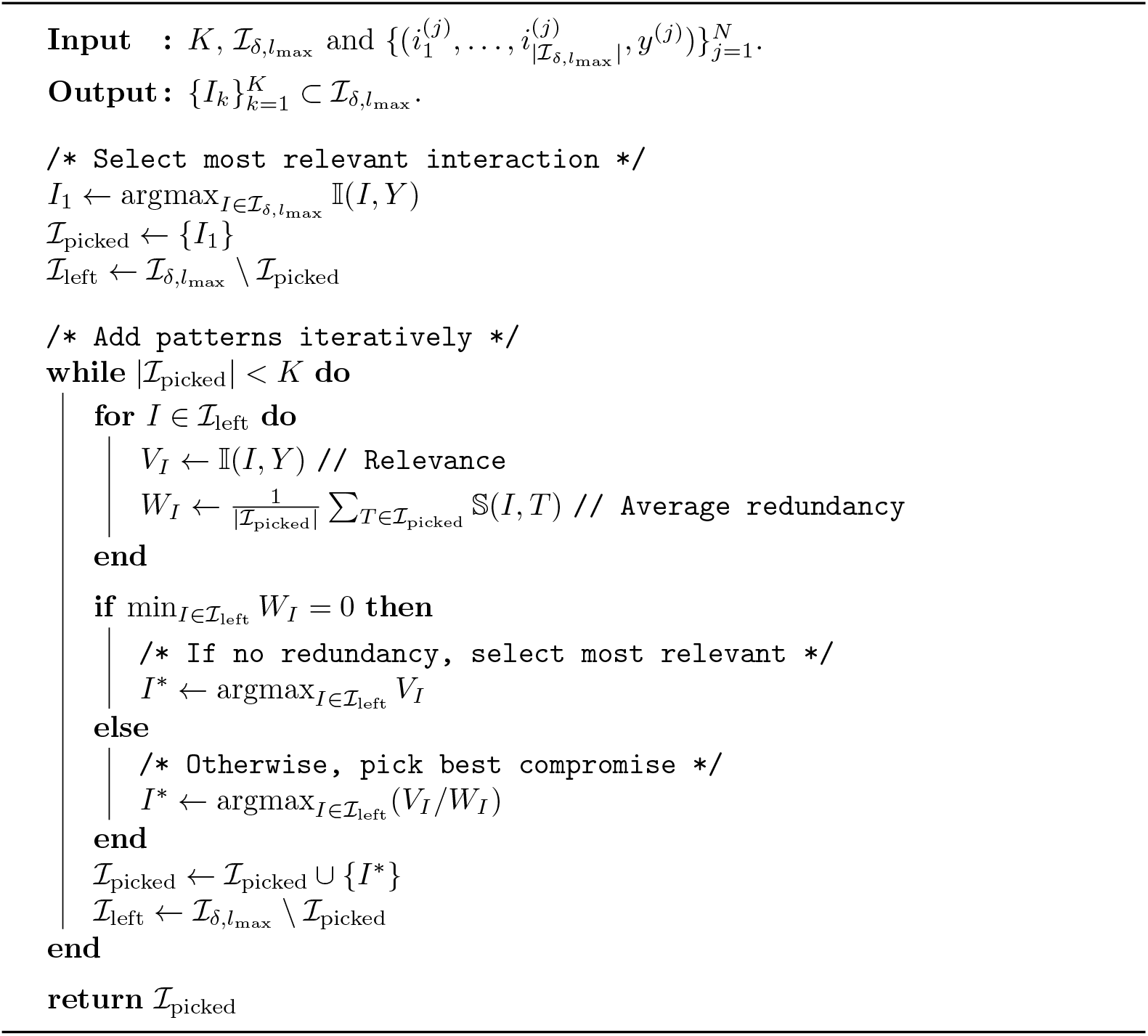

We remark that *K* is an hyperparameter chosen by the user. In particular, one may as well optimize its value accordingly to some grid-search algorithm of choice. Indeed, the most computationally expensive parts in the *hi*PRS pipeline are the candidates search and the computation of relevance/redundancy measures. Once these steps have been carried out, multiple values of *K* can be tested and the user may choose the one considered to be optimal.

Once the set 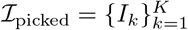 has been identified, we compute the weights *β*_0_, …, *β*_*K*_ by fitting the Logistic Regression model below

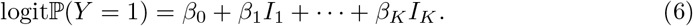

Finally, we define the *hi*PRS accordingly to (3), meaning that the score is actually the (affine) linear predictor in (6).

### Benchmark Algorithms

We report below a short description and the benchmark algorithms together with their implementation details.

#### Penalized PRSs with additive effects

From this class of methods we chose three of *hi*PRS competitors, namely Lasso [31], Ridge [32] and Elastic-Net [33]. These algorithms are traditional penalized LR models that only account for the additive main effect of the predictors on the target variable, while imposing an *𝓁*_2_ bound of the form 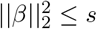 (Ridge) or an *𝓁*_1_ bound ||*β*|| *≤ s* (Lasso), or the combination of the two (Elastic-Net) on the coefficients. Note that *𝓁*_1_ penalizations also perform feature selection by shrinking coefficients to zero, making these approaches popular to model polygenic risk from large genotype data. Fundamentally, the penalization terms enter the loss function during the training phase, and their contribution is weighted by some hyperparameter, e.g. *λ* for Lasso and Ridge, *λ*_1_ and *λ*_2_ for Elastic-Net. To run these algorithms in our experiments, we relied on the implementation available in the Python library scikit-learn, with default values for *λ, λ*_1_ and *λ*_2_. All the code was written in Python 3.7.

#### Glinternet

Glinternet is a method for learning pairwise interactions in a LR satisfying strong hierarchy: whenever an interaction is estimated to be nonzero, both its associated main effects are also included in the model. The idea of the algorithm is based on a variant of Lasso, namely group-Lasso [46], that sets groups of predictors to zero. glinternet sets up main effects and first-order (i.e., pairwise) interactions via groups of variables and then intuitively selects those that have a strong overall contribution from all their levels toward explaining the response. For more formal definitions we refer the reader to [34].

The amount of regularization is controlled by *λ*, with larger values corresponding to stronger shrinkage and less interactions included. The freely distributed R implementation of glinternet allows the user to define the number of pairwise interactions (*n_int*) to find and the size of a grid of *λ* values of decreasing strength to fit the model with. This grid is built automatically by splitting equally from *λ*_*max*_, that is data-derived as the value for which all coefficients are zero, to the minimum value, computed as *λ*_*min*_ = *λ*_*max*_*/*0.01. The algorithm will fit this path of values and stop when *n_int* is reached. For this experiment we set *n_int* = [3, 8], defining a grid of 100 *λ*s. All other hyperparameters were left to default settings.

#### DNN-Badre

DNN-Badre [36] constructs a Deep Feed Forward Neural Network (DNN) to predict a binary outcome. To implement their approach, we built a DNN architecture with the same specifics provided by the authors, i.e. number of layers, neurons and activation functions. We trained the models via stochastic gradient descent, with mini-batches of batch size 10 and for a total of 200 epochs. The optimization was carried out using the Adam optimizer, with a learning rate of 10^*−*3^. All the code was written in Python 3.7, mostly using the Pytorch library.

#### SVM-Behravan

SVM-Behravan is a multi-step ML-based algorithm recently presented as a state-of-the-art approach to polygenic risk scoring [35, 47]. We will provide here an intuitive description of the algorithm with the needed details to understand our settings for the present work, while for more technical specifications we refer the reader to the seminal work in [35]. The algorithm presented in Behravan et al. (2018) is composed of a so called *first module*, where an XGBoost is used to evaluate the importance of SNPs on a risk prediction task by providing an initial list of candidate predictive SNPs. The authors use the average of feature importances (a.k.a. “gain”) provided by the gradient tree boosting method, as the contribution of each SNP to the risk. Then, in the *second module*, the candidate SNPs are used for an adaptive iterative search to capture the optimal ways of combining candidate SNPs to achieve high risk prediction accuracy on validation data. In particular, top *M* and bottom *M* SNPs from the candidate list are ranked separately based on accuracy, then top and bottom *N* SNPs are switched between the two lists. The process is repeated with *M* (i.e., *window size*) increased of *W* at each iteration, until the two sublists overlap and the optimal ranking is achieved. Finally, an SVM is trained to distinguish cases (positive samples) and controls (negative samples) using the *S* top-ranked SNPs in the optimal ranking as feature vectors and a linear kernel. In the original paper the performance of the algorithm is averaged across 5-fold CV, meaning that the pipeline from first module to SVM is repeated 5 times on different folds of the training and test set. As the author included this step to overcome the problem of small samples to train high-performance risk prediction models, we followed their instructions and for each of the 30 simulated dataset we performed the 5-fold CV. This lead to the 5 different subsets of SNPs selected as shown in the right panel of Fig. 5. However, to keep the running time of this computationally very expensive method reasonable and for a fair comparison with all the other methods for which we did not perform extensive optimal hyperparameter search, we avoided the very long optimization of the XGBoost model by setting manually the optimal hyperparameters described in the paper. In particular, we set *M* = 2, *W* = 1 and *N* = 1.

### Performance Metrics

In imbalanced settings, the Accuracy of a classifier (i.e., the fraction of correctly classified observations) can be a miselading metric to assess a model performance. Therefore, to evaluate *hi*PRS and the benchmark algorithms in predicting the outcome, we exploited two metrics that combined provide a full picture of the real performance of these methods, expecially in predicting the most interesting class (i.e. the positive class), which is oftentimes the underrepresented one in real data. Therefore, for our experiments we chose the Area Under the receiving operating characteristic Curve (AUC) and the Average Precision (AP).

The two are computed as follows. For any fixed discrimination threshold, let *TP* and *FP* be the true and false positives, respectively. Similarly, let *TN* and *FN* be the true and false negatives, respectively. First, we derive the following quantities,

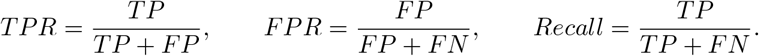

that is, the True Positive Rate (TPR), the False Positive Rate (FPR) and the Recall. TPR is the fraction of true positives out of the positives, also known as *sensitivity* or *precision*. FPR, or *specificity*, is the fraction of false positives out of the negatives. Finally, the Recall quantifies the ability of the classifier to recognize positive samples. By plotting *TPR* against *FPR* for various discrimination threshold levels, we obtain the ROC curve. Conversely, the pair (*Recall, TPR*) yields the precision-recall curve. AUC summarizes the performance of a binary classifier by computing the area under the ROC curve, while AP considers the area under the precision-recall curve. In practice, the two areas are estimated using quadrature rules. In our implementation, we employ the trapezoidal rule for AUC, while we use the rectangular rule for AP.

## Data Availability Statement

The DACHS data that support the findings of this study are available on request. The data are not publicly available due to privacy or ethical restrictions. The code developed to reproduce the experiments carried out in this work, and to apply the proposed *hi*PRS to any other data, is freely distributed on a GitHub repository that can be accessed through this link: github.com/NicolaRFranco/hiprs

## Funding

This project has received funding under the ERA-NET ERA PerMed / FRRB grant agreement No ERAPERMED2018-244, RADprecise - Personalized radiotherapy: incorporating cellular response to irradiation in personalized treatment planning to minimize radiation toxicity. DACHS study was supported by the German Research Council (BR 1704/6-1, BR 1704/6-3, BR 1704/6-4, BR 1704/6-6, CH 117/1-1, HO 5117/2-1, HE 5998/2-1, KL 2354/3-1, RO 2270/8-1, BR 1704/17-1), the Interdisciplinary Research Program of the National Center for Tumor Diseases (NCT), Germany, and the German Federal Ministry of Education and Research (01KH0404, 01ER0814, 01ER0815, 01ER1505A, 01ER1505B, 01GL1712).

## Notes

### Competing Interest Statement

The authors have declared no competing interest.

